# Transposable elements are constantly exchanged by horizontal transfer reshaping mosquito genomes

**DOI:** 10.1101/2020.06.23.166744

**Authors:** Elverson Soares de Melo, Gabriel da Luz Wallau

## Abstract

Transposable elements (TEs) are a set of mobile elements within a genome. Due to their complexity, an in-depth TE characterization is only available for a handful of model organisms. In the present study, we performed a *de novo* and homology-based characterization of TEs in the genomes of 24 mosquito species and investigated their mode of inheritance. More than 40% of the genome of *Aedes aegypti, Aedes albopictus*, and *Culex quinquefasciatus* is composed of TEs, varying substantially among *Anopheles* species (0.13%–19.55%). Class I TEs are the most abundant among mosquitoes and at least 24 TE superfamilies were found. Interestingly, TEs have been continuously exchanged by horizontal transfer (212 TE families of 18 different superfamilies) among mosquitoes since 30 million years ago, representing around 6% of the genome in *Aedes* genomes and a small fraction in *Anopheles* genomes. Most of these horizontally transferred TEs are from the three ubiquitous LTR superfamilies: Gypsy, Bel-Pao and Copia. Searching more 32,000 genomes, we also uncover transfers between mosquitoes and two different Phyla—Cnidaria and Nematoda—and two subphyla—Chelicerata and Crustacea, identifying a vector, the worm *Wuchereria bancrofti*, that enabled the horizontal spread of a Tc1-mariner element of *irritans* subfamily among various *Anopheles* species. These data also allowed us to reconstruct the horizontal transfer network of this TE involving more than 40 species. In summary, our results suggest that TEs are constantly exchanged by common phenomena of horizontal transfers among mosquitoes, influencing genome variation and contributing to genome size expansion.

**Author Summary:** Most eukaryotes have DNA fragments inside their genome that can multiply by inserting themselves in other regions of the genome, generating variability. These fragments are called Transposable Elements (TEs). Since they are a constituent part of the eukaryote genomes, these pieces of DNA are usually inherited vertically by the offspring. To avoid damage to the genome caused by the replication and insertion of TEs, organisms usually control them, leading to their inactivation. However, TEs sometimes get out of control and invade other species through a horizontal transfer mechanism. This dynamic is not known in mosquitoes, a group of organisms that acts as vectors of many human diseases. We collected mosquito genomes available in public databases and characterized the whole content of TEs. Using a statistic supported method, we investigate TE relations among mosquitoes and discover that horizontal transfers of transposons are common and occurred in the last 30 million years among these species. Although not as common as transfers among closely related species, transposon transfer to distant species also occur. We also identify a parasite, a filarial worm, that may have facilitated the transfer of TE to many mosquitoes. Together, horizontally transferred TEs contribute to increasing mosquito genome size and variation.

## 1. Introduction

Transposable elements (TEs) are DNA sequences that can move from one genome locus to another. They were discovered by Barbara McClintock in the late 1940s, in the maize genome, but it was not until the 1970s that they were rediscovered in other species and their genetics and evolutionary importance revealed in all branches of life [1,2]. It is now known that TEs profoundly reshape the genomes of all species in a myriad of ways. The repetitive nature of TEs generates abundant sites for chromosomal rearrangements and their transposition may have serious consequences for the host genomes by way of insertion into host gene coding regions or introns. This may give rise to new poly(A) tails, causing exon skipping or creating alternative promoters [3,4]. Although TEs are part of the host genome, they can replicate, generating new copies of itself, using the host’s molecular machinery, thereby triggering a coevolution arms race. Such intertwined interaction may have wide-ranging consequences for the organisms’ evolution [5–7].

There is a large diversity in sequence-structure among TEs. Classification schemes distinguish two large groups of TEs based on the intermediate transposition molecule: TEs that are mobilized via an RNA intermediate or retrotransposons and TEs that transpose via a DNA intermediate or DNA transposons [8,9]. Many retrotransposons are large elements ranging from 10 to 20kb in size showing a number of domains and *long terminal repeats* (LTRs) which are also normally found in viral genomes hence sharing a common origin with different viral taxa. Another fraction of retrotransposons is composed of highly diversified elements that usually have poly(A) tail and an endonuclease or apurinic endonuclease, usually known as non-LTR retrotransposons [10]. DNA transposons are divided into two subclasses distinguished by transposition mechanism: subclass I, are small elements (2 to 7 kb) normally coding for a single protein comprising elements that transpose by cleaving both strands of DNA, and subclass II is composed of large (Mavericks reaching 40 Kb) and medium-sized TEs (Helitrons reaching 15 Kb) that transpose with cleavage of only one DNA strand [8,11,12].

TEs are inherited by vertical transfer, the transfer of genetic from ancestral to descendant species, to all host offsprings [13]. However, there is growing evidence that TEs can also move horizontally between independent species by a phenomenon known as horizontal transfer (HT) [14]. Recent large-scale studies on Insects and Vertebrates have revealed thousands of Horizontal Transposon Transfer (HTT) events and general patterns, such as the occurrence of HTT more frequently between closely related host species which share a spatiotemporal overlap; the substantial contribution of horizontally transferred TEs to the genome size of the host species; and that DNA transposons are transferred horizontally much more frequently than Retrotransposons [15]. However, many open questions need further attention to understand HTT at different host taxa, its functional impact on host genomes, and by which mechanisms HTT occurs. In a simplistic view, HTT might occur by direct transfer of a TE between species or mediated by a vector/intermediate species that transport TEs between hosts [16]. Currently, there is evidence scattered on different host/TE systems that these elements can ‘hitch a ride’ on other parasite genomes/particles, such as those of parasitic nematodes, macroscopic blood-sucking insects, and viruses that interact intimately with different host species [17–19]. However, the exact mechanisms by which HTT occurs has remained as a large mystery since it is difficult to reproduce such phenomenon in the lab. Moreover, reconstruct TEs horizontal steps from sequence data is full of pitfalls and missing information mainly due to incomplete sampling of host-intermediate species [16,20].

TEs make up a significant portion of the genome of some species: 80% of the model plant organism *Zea mays* [21], and almost half of the genome of the mosquito *Aedes aegypti* [22]. Mosquito belongs to Culicidae family, which comprises more than 3500 known species, dispersed trough all continents except Antarctica. It is divided into two large subfamilies, Culicinae and Anophelinae, the former is composed of 11 tribes with several genera, while the latter has only 3 genera, having the *Anopheles* genus as the most species-rich group [23]. Many mosquito species of the Culicidae family are important human pathogen vectors of dengue, chikungunya, Zika, and yellow fever viruses as well as other pathogens, such as unicellular eukaryotes of the *Plasmodium* genus and filarial nematodes such as *Wulchereria bancrofti* and *Brugia malayi* [24]. In order to devise new methodologies to control such pests, at least 24 mosquito genomes have been sequenced from three Culicidae genera: *Aedes, Anopheles*, and C*ulex* [22,25–29]. TE content varies substantially in mosquito genomes, with a greater abundance of Class I over Class II TEs [22,25], but well-characterized mobilomes are restricted to the most extensively studied species, such as *An. gambiae* and *Ae. aegypti* and even for those species there is no TE data accessible or standardized to allow more in-depth comparative analysis [30]. Therefore, there is little research regarding the inheritance mechanism of TEs among mosquitoes and only four well-documented horizontal transfer cases have been reported so far [31].

In this study, we recharacterized the TE content of 24 mosquito species and evaluated the horizontal spread of these TEs inside and outside of the Culicidae family. As result, we found hundreds of horizontal TE transfers that occurred in the past 30 million years among these 24 species and observed that some of these are involved in HTTs with distantly related species from other Phyla. We uncovered a vector species, the nematode *Wulchereria bancrofti*, that facilitated the horizontal spread of a specific TE among mosquitoes. Besides, horizontally transferred TEs contributed significantly to the genomic expansion of mosquito genomes and we found an absence of positive correlation between HTT events and spatiotemporal overlap. Overall, our data brought a substantial contribution to understanding the HTT phenomenon and its impact on host genomes.

## 2. Results

### 2.1 TE content in mosquito genomes

We performed a recharacterization of TEs in 24 species of mosquito using a *de novo* and a homology-based approach. A substantial variation was detected in the TE content of several species, for instance, *An. gambiae*—the most extensively studied mosquito species and the one that has the largest number of characterized TEs stored in databanks showed a TE content of 20.23% by TEdenovo, 14.94% by homology search against Repbase and 13.35% against TEfam (Fig 1A). The discrepancy between the three methods is more pronounced in one of the least studied species, *An. darlingi*. A 10-fold difference was observed between TEdenovo (1.97%) and homology-based search against Repbase (0.19%) (Fig 1A). To generate a final dataset of the TEs recovered using each methodology, we used all TEs recovered by way of three different approaches and masked them using RepeatMasker. The final proportion of TEs in each genome is shown in S1 Fig.

**Fig 1:**
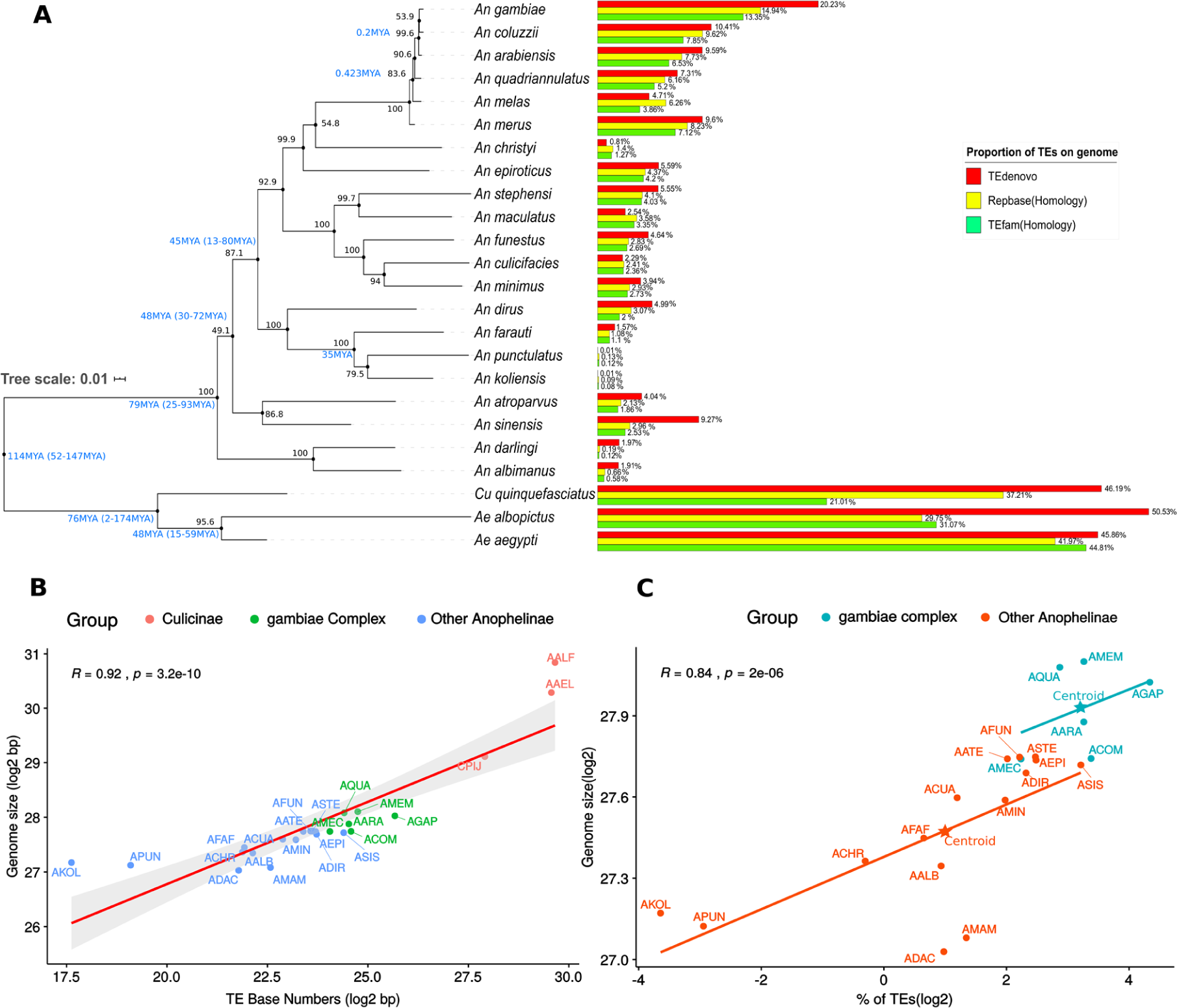
Transposable elements of mosquitoes. (A) A maximum likelihood-based phylogenetic inference, with bootstrap support over the nodes (see Materials and Methods - Genomes Analyzed), reporting the proportion of genome size occupied by transposable elements; blue letters values near bootstrap support represent medium divergence time between species obtained from TimeTree [95], colored horizontal bars represent the proportion of TEs characterized by different methods. (B) Correlation analysis of mosquito genome sizes and TE fraction. In the *An. gambiae* species complex, non-TE elements make a greater contribution to genome size than in other Anophelinae (C), for the meaning of four-letter species abbreviations see Table 1.

The An. *gambiae* complex, which contains six African species--*An. gambiae, An. coluzzii, An. arabiensis, An. quadriannulatus, An. melas*, and *An. merus*-- showed the highest proportion of genome covered by mobile elements of all *Anopheles* species investigated. At the other extreme, the New Guinean mosquitoes *An. farauti, An. punctulatus*, and *An. koliensis* showed a very low TE content of between 0.13% and 1.57%, considering the highest estimates for all methodologies. Since neither a homology-based search nor the *de novo* approach identified a TE content greater than 1% in the last two species, we tried other methods of TE characterization using raw sequencing reads to ascertain whether these estimates might have been biased by TE removal steps during the original genome assembly. However, these small TE proportions were confirmed (data not shown). On the other hand, the three mosquito species of the Culicinae subfamily (*Ae. aegypti, Ae. albopictus*, and C*ulex quinquefasciatus*) have a much higher TE content than Anophelinae species, ranging from around 44% to more than 50% in *Ae. aegypti* and *Ae. albopictus* respectively (S1 Fig).

**Table 1.** Number of TE families in each species that is involved in HTT events

| Species | Code | Class II | Class I |  | Total |
| --- | --- | --- | --- | --- | --- |
|  |  |  | Non-LTR | LTR |  |
| *Anopheles merus* | AMEM | 14 | 15 | 26 | 55 |
| *Aedes aegypti* | AAEL | 12 | 13 | 27 | 52 |
| *Aedes albopictus* | AALF | 10 | 14 | 27 | 51 |
| *Anopheles gambiae* | AGAP | 14 | 6 | 29 | 49 |
| *Anopheles epiroticus* | AEPI | 7 | 15 | 16 | 38 |
| *Anopheles atroparvus* | AATE | 10 | 7 | 18 | 35 |
| *Anopheles dirus* | ADIR | 9 | 9 | 11 | 29 |
| *Anopheles stephensi* | ASTE | 5 | 8 | 15 | 28 |
| *Anopheles maculatus* | AMAM | 6 | 9 | 12 | 27 |
| *Anopheles minimus* | AMIN | 7 | 9 | 11 | 27 |
| *Anopheles sinensis* | ASIS | 10 | 6 | 9 | 25 |
| *Anopheles funestus* | AFUN | 8 | 1 | 12 | 21 |
| *Anopheles melas* | AMEC | 11 | 0 | 7 | 18 |
| *Anopheles arabiensis* | AARA | 5 | 3 | 8 | 16 |
| *Anopheles quadriannulatus* | AQUA | 7 | 3 | 5 | 15 |
| *Anopheles culicifacies* | ACUA | 5 | 3 | 4 | 12 |
| *Anopheles coluzzii* | ACOM | 3 | 0 | 4 | 7 |
| *Anopheles farauti* | AFAF | 0 | 5 | 0 | 5 |
| *Anopheles albimanus* | AALB | 0 | 2 | 0 | 2 |
| *Anopheles christyi* | ACHR | 0 | 0 | 2 | 2 |
| *Anopheles darlingi* | ADAC | 0 | 0 | 0 | 0 |
| *Anopheles koliensis* | AKOL | 0 | 0 | 0 | 0 |
| *Anopheles punctulatus* | APUN | 0 | 0 | 0 | 0 |
| *Culex quinquefasciatus* | CPIJ | 0 | 0 | 0 | 0 |

The final TE content obtained from masking all TEs recovered was used to evaluate whether the mosquito’s genome size correlates with the mobilome content. The analysis indicated a strong positive correlation (R=0.92) between TE content and mosquito genome size (Fig 1B). Looking only to *Anopheles* species there are two patterns, species of the *An. gambiae* complex have a larger genome size for similar TE content compared to other Anophelinae mosquitoes (Fig 1C), suggesting that TEs play an important role in driving genome size in the *An. gambiae* complex. However, other phenomena at the genomic scale might also contribute to genome expansion while TE driven expansion has a more significant role in driving genome size in the remaining *Anopheles* species.

### 2.2 Distribution of superfamilies by mosquitoes

At least 24 TE superfamilies were found in the mosquito genomes (Fig 2). This diverse mobilome comprises most of the transposable element orders established by Wicker et al. (2007) and some superfamilies that had not been described at that time (S2 Fig). LTR superfamilies, such as Copia, Gypsy, and Bel-Pao, are ubiquitous in mosquito genomes (Fig 2), accounting for at least 6% of the *Ae. aegypti* genome and 4.3% of *An. gambiae*. LINE elements are abundant in most species, they represent 14.5% of the genome of *Ae. aegypti*, 15.7% of that of *Ae. albopictus*, and 6.2% of that of *Cu. quinquefasciatus*. RTE and Jockey are the most abundant superfamilies in mosquitos’ genomes, while R2 is sparse. Twelve different TIR superfamilies are well represented in mosquitoes. As expected, Tc1-Mariner—the most widespread superfamily of TEs among arthropods—is present in all species of mosquitoes, even those with a small TE content. Non-autonomous TIR elements, such as putative MITEs and SINEs, are also present in the majority of species. It was not possible to classify some elements at the superfamily level, most of these have been classified only as Class I and Class II (S2 Fig). These elements represent around 15 - 20 % of all TEs in mosquitoes of *An. gambiae* species complex and 18% of the *Cu. quinquefasciatus* genome, but represent only a small proportion of other mosquito genomes (2E^-4^ to 3 %).

**Fig 2:**
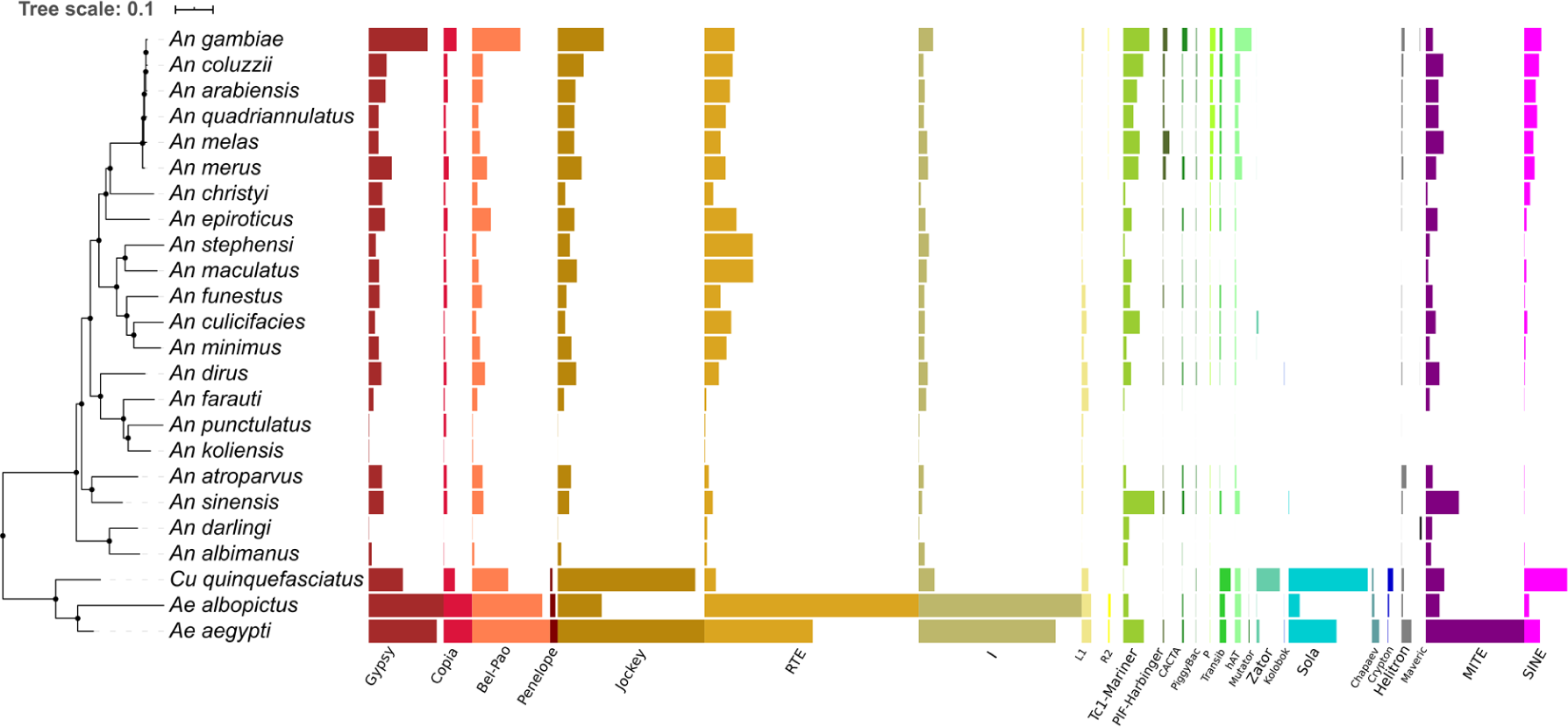
Proportion of each TE superfamily in the genomes of mosquitoes. A phylogenetic tree of 24 mosquito species showing the distribution of different TE superfamilies in mosquitoes. Each colored bar represents the proportion of different TE families from the same superfamily (names in X-axis) following Wicker et al. (2007) hierarchical scheme. Uncharacterized or order-level-characterized TEs not shown (See S2 Fig).

Chimeric elements are formed by elements from two or more distinct superfamilies or have at least one characteristic that derives from more than one superfamily. This type of element is present in almost all species, but a higher proportion of chimeric TEs was found in the *Ae. aegypti, An. arabiensis, An. gambiae*, and *An. quadriannulatus* genomes (S2 Fig).

Both Class I and II superfamilies are heterogeneously distributed among mosquitoes. The Penelope superfamily was only found in the three Culicinae mosquitoes studied and covered only a small fraction of these genomes (Fig 2). The same pattern can be seen with Mutator, Chapaev, and Crypton-like elements. The most remarkable difference in superfamily distribution was found in the case of Sola elements. This element is one of the most abundant TEs in the *Cu. quinquefasciatus* genome (2.86%). It is also present in *Aedes* mosquitoes and only one Anophelinae species—*An. sinensis* (representing nearly 0.01% of its genome). Another clear difference is the absence of elements from the CACTA, P, and Transib superfamilies among neotropical and New Guinean anophelines. R2 is another superfamily that has an interesting distribution. It is present only in *Aedes* mosquitoes and in species from the *An. gambiae* complex (Fig 2).

From the host mosquitos’ genomes perspective, the *An. gambiae* species complex has a similar TE superfamily landscape, while there is substantial variation in distribution and abundance between *Aedes* and *Anopheles* mosquitoes (Fig 2). However, it is clear that *Cu. quinquefasciatus* has the most distinct TE superfamily landscape of the Culicidae family, with the greatest abundance of Zator, Sola, and Crypton-like elements compared to all other mosquito species studied. Another distinctive feature of the *Cu. quinquefasciatus* mobilome is the low genome proportion of ubiquitous superfamilies among mosquitoes such as Tc1-Mariner, RTE, and I (Fig 2).

### 2.3 Relative TE ages per mosquito species

To examine the landscape of TEs within species genomes over time, we estimated the K2P distance of all TEs. We clustered TEs in the four highest abundant orders: LTR, LINE, TIR, and SINE. The boxplots in S3 Fig reveals interesting patterns: mosquitoes from the *An. gambiae* complex tend to have more recent TE families from the TIR, LINE, and LTR orders than other Anophelinae. Low K2P values in these six mosquitoes indicate that many TE families are currently active or had been recently active. In contrast, the vast majority of LINE families in *An. darlingi, An. albimanus, An. punctulatus* and *An. koliensis*—which are species with a very low TE load—are old and probably no longer active.

Most SINE elements in the genome of *Anopheles* mosquitoes are middle-aged (S3 D Fig) and seems to have undergone ancient transposition bursts and no longer appear to be active in these mosquitoes. A distinct pattern can be seen in Culicinae mosquitoes, where we detected recent expansion of SINEs. In general, the K2P pattern provides clues that TEs are younger and more recently active in Culicinae than Anophelinae families.

### 2.4 Hundreds of HTT events have occurred among the mosquito species

We found 212 TE families with significant HT signals among the species studied. However, it is important to note that this is certainly an underestimate of the true number of HTT events since most TE families have more than one significant pairwise HTT signal (Fig 3A). We, therefore, decided to describe it in more general terms, retaining at least one HT event for each TE family, in view of the complexity of the signals found and the lack of currently available algorithms for determining the most likely HTT scenario and estimating the minimum number of events required to explain the signal.

**Fig 3:**
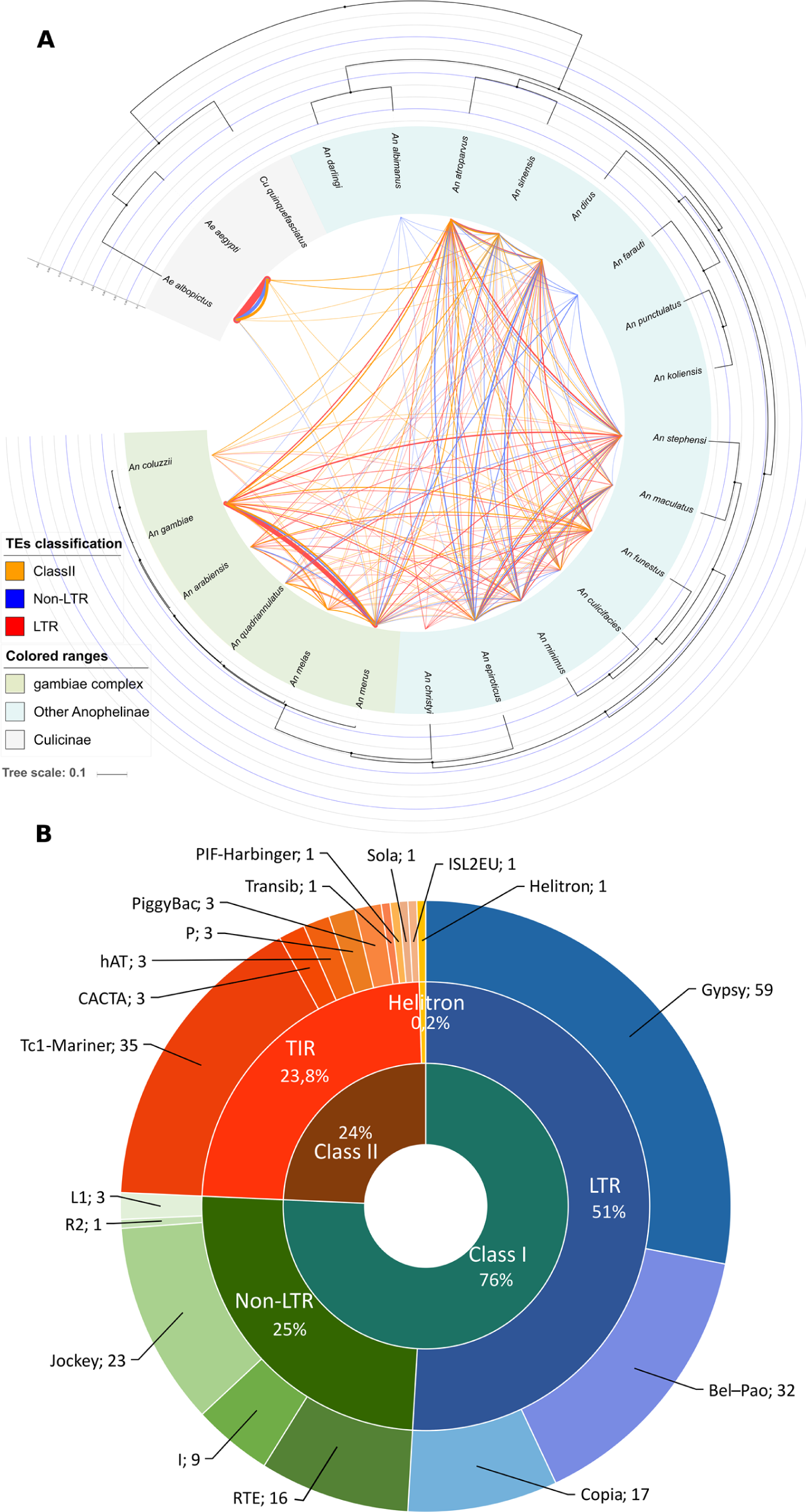
HTT among mosquitoes and representative TE superfamilies. (A) A phylogenetic tree (see Materials and Methods - Genomes Analyzed) whose edges represent pairwise HTT events and thickness is proportional to the number of events. (B) Classification of these 212 TE families involved in HTT among mosquitoes.

More than three-quarters of transposons involved in HT events belong to Class I TEs, covering eight different superfamilies (Fig 3B). The superfamily with the largest number of families undergoing HT is Gypsy, in which HT accounts for more than 35% of all events, followed by Tc1-Mariner elements. Although Class I TEs are more abundant in mosquito genomes and cover most TE families involved in horizontal transfer events, Class II TEs were found to contain more superfamilies (9) undergoing HT. In this group, the Tc1-Mariner superfamily deserves special mention, since it has 11 TEs involved in HT events involving four or more mosquito species.

Almost 25% of the HT signals detected (47 families) occurred only between *Ae. aegypti* and *Ae. albopictus*. One hundred and sixty-one TE families involved in HT events were detected among species of the *Anopheles* genus, accounting for 18 of the 21 species. Only tree mosquito species, *An. darlingi, An. punctulatus* and *An. koliensis*, were not involved in HTT events. One *Anopheles* species, *An. merus*—an early divergent species of the *A. gambiae* complex—was involved in the largest number of HTTs (Fig 3A). The majority of species from the *Anopheles* genus are involved in HTTs from both Class I and Class II TEs (Table 1). In three species we detected only HTTs of Class I TEs: non-LTR elements in *An. albimanus* and *An. farauti*, and LTRs elements in *An. christyi*. Concerning HTT between different mosquito genera, six events were found. Five out of six of those belong to the Tc1-Mariner superfamily, while only one R4—a non-LTR retrotransposon —was transferred from *Ae. albopictus* to the ancestor of *An. gambiae* complex mosquitoes, as already described in a previous study [32]. No horizontal transfer events involving the *Cu. quinquefasciatus* species were found. It is important to note that all HTT events showed very low p-values (S4 Fig) associated with several additional HTT evidence as patchy distribution and phylogenetic incongruences between TE and host phylogeny, as further discussed below (**2**.**8 - Horizontal transfer of TEs involving distantly related eukaryotic species**).

We estimated the HTT rate for Class II, LTR, and non-LTR elements based on the number of TE families with HT signal on the total number of families tested with VHICA. Class II TEs showed the highest rate (24.1%), followed by non-LTRs (20.2%) and LTRs (19.4%). Around 14% of the HTT networks involve at least four species. This proportion is small, particularly among LTR families, where four or more mosquito species were detected in less than 10% of the total number of LTR families involved in HTT. Of all the superfamilies studied, Tc1-Mariner and RTE are the ones whose elements are most often horizontally transferred across many species, 17 and 13 respectively. Details about the participation of species in each TE family horizontal transfer network can be seen in Supplementary File 2, which lists all VHICA output images of positive HTT cases. The relative age of each TE family undergoing HT per mosquito genome is given in S5 Fig.

### 2.5 The direct impact of HTT on mosquito genomes

TEs horizontally transferred between species represent a significant fraction of some mosquito genomes (Fig 4A). We estimated that about 6% of *Ae. aegypti* and *Ae. albopictus* genomes are covered by horizontally transferred TEs representing more than 10% of the total TE content of these species. In the *Anopheles* genus, the impact of horizontal transfer on genome size is much smaller. Only *An. gambiae* and *An. coluzzii* showed more than 1% of their genome covered with copies derived from horizontally transferred TEs.

**Fig 4:**
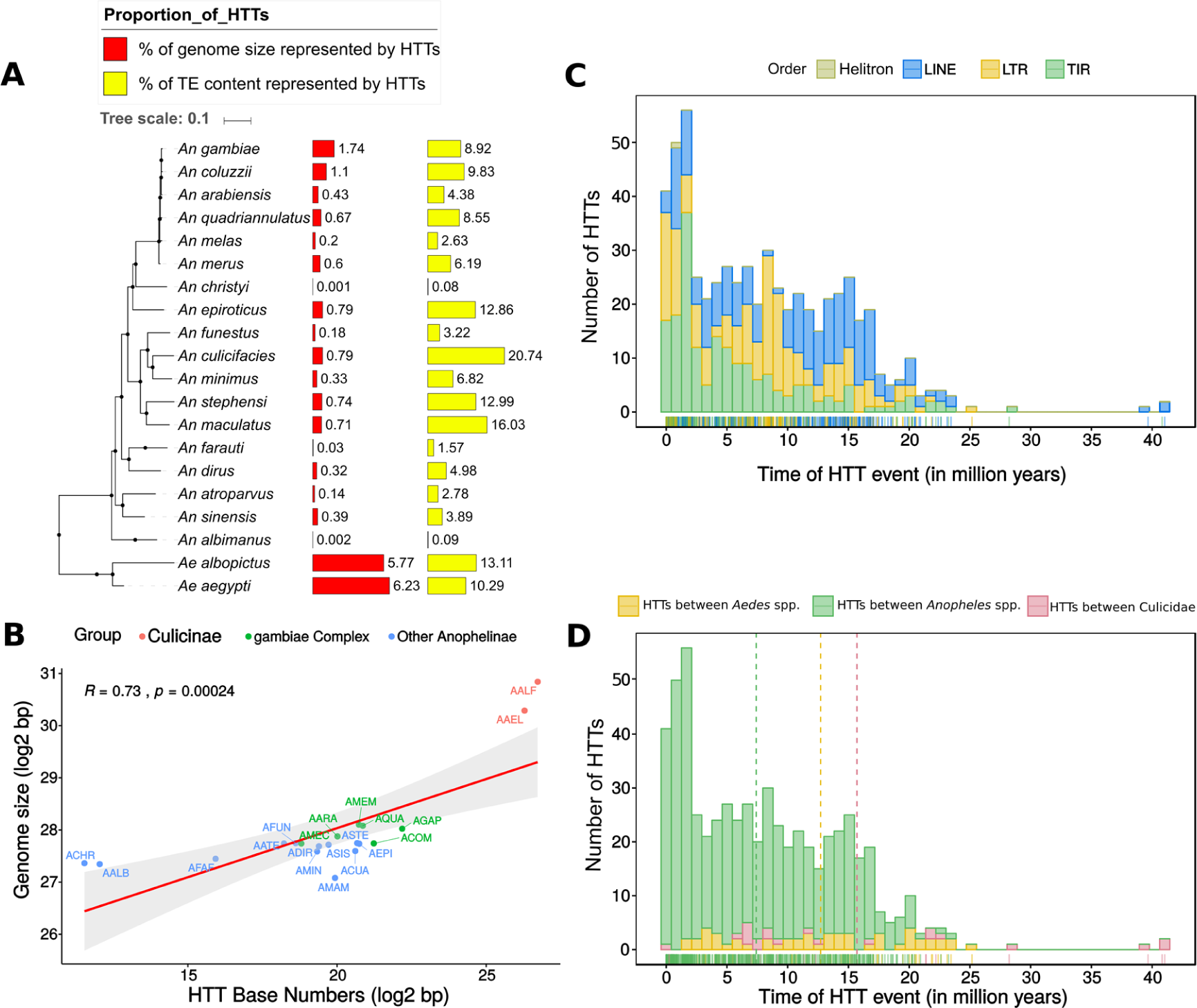
Influence of HTT on the host genome and dating. (A) The proportion of the host genomes composed of TEs involved in HT. (B) The correlation between the number of nucleotides covered by TEs involved in HTT events and genome size in mosquitoes. The age of each pairwise HTT event estimated and represented by TE groups (B) and host taxa (C).

The proportion of genome covered by horizontally transferred TEs showed a positive correlation with genome size of the host species Fig 4B. Therefore, TE invasion by way of horizontal transfer may significantly contribute to genomic expansion and also impact the size of the mosquito genome. Thus, the more frequent HTT events, the larger the genome.

### 2.6 Horizontal transfer of transposable elements occurred in the last 30 million years

We estimated the HTT dating based on all pairwise 584 significant comparisons found. The vast majority of HTT events detected occurred in the last 30 million years (Figs 4C and 4D), a time during which most *Anopheles* species underwent speciation [33]. Only one transfer is older than this: a transfer of an R4 clade involving species from the *An. gambiae* complex and both of the *Aedes* species studied that took place around 40 million years ago (S6 Fig). Non-LTRs have the most uniform number of HTTs over time, while HTT among LTRs and TIR elements intensified more recently (Fig 4C). In general, horizontal transfers occurring between species of the *Anopheles* genus are more recent than those taking place within the *Aedes* genus or between these two genera. Most of the HTTs between *Ae. aegypti* and *Ae. albopictus* identified are relatively ancient transfers, most having occurred more than 15 million years ago (Fig 4D).

### 2.7 Geographical location does not affect the number of transfers between mosquito species

In order to investigate whether geographical overlap of mosquito distribution range favors the occurrence of HTTs, we compared the distribution of species across the globe with the number of HTTs between each pair of species. No correlation was found between these variables. (R = 0.03, p-value = 0.73) (S7 Fig).

### 2.8 Horizontal transfer of TEs involving distantly related eukaryotic species

We also examined whether some of the transposable elements that are horizontally transferred among mosquitoes might also be involved in transfers to other eukaryotes. Five TE families from the *Tc1-Mariner* superfamily showed a wider horizontal transfer network with other species. Three of them have already been described in previous studies, showing transfers between at least two species (Fig 5). Our results further expand this network, adding twenty new species. The other HTT events in the remaining two TE families are described here for the first time. The similarity between mosquito elements and those of other eukaryotic species can be seen in Supplementary File 3.

**Fig 5:**
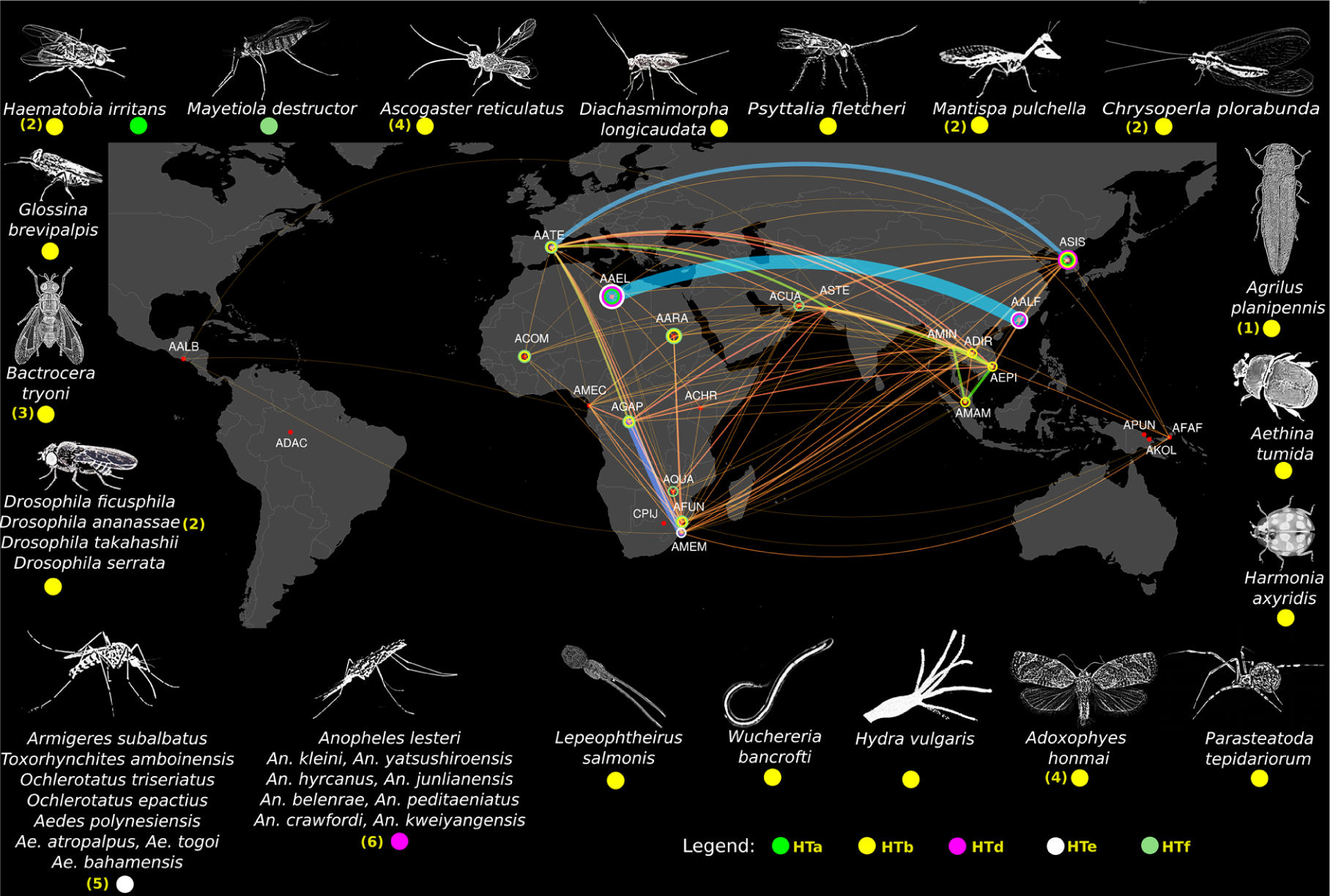
Mosquito HTT network. Red dots represent the species sampling site for the genomes used in this study. The lines connecting dots represent significant pairwise HTT events. Thickness and colors of the edges are proportional to the number of HTT events. Five TEs are horizontally transmitted to other species outside the Culicidae family. These species are shown on the edge of the map. Colored circles represent the five TE groups: HTa, b, d, e, and f. The colored circle around red dots indicates that this mosquito species is involved in a long-range HTT network of at least one specific TE (HTa to HTf). The numbers inside the round brackets represent some preliminary evidence of HTTs involving these TEs and species in other published articles. 1 represents the APmar1 element already described by Rivera-Vega et al. [96], 2 represents HTT transfers among species described by Robertson and Lampe [97], 3 represents an element described by Green and Frommer [98], 4 represents HTT transfer between *Ascogaster reticulatus* and *Adoxophies honmai*, 5 represents HTT events already described in mosquitoes [99], 6 represents HTT events already described in other *Anophelinae* species [100] not investigated in our study.

The species involved in HTT events belong to three metazoan phyla. The first phylum was Arthropoda, which covers the majority of species. It is represented by the class Arachnida, by the common house spider *Parasteatoda tepidariorum*, by Hexanauplia, represented by the copepod *Lepeophtheirus salmonis*, and by the class Insecta, which includes most of the mosquito species involved in HTT. The second phylum was Cnidaria, with only one representative, *Hydra vulgaris*. The third was Nematoda, where we found a horizontal transfer case involving the parasitic worm *Wuchereria bancrofti* and other *Anopheles* species that transmit this nematode to humans. Besides the high identity between mosquito TEs and those of distantly related species, sometimes separated by more than 700 million years, we also observed a patchy distribution of these elements, providing further support for the horizontal transfer events.

Two of these TE lineages (HTa and HTe) belong to the ITmD37E group, which includes TEs that have the DD37E motif on their transposase sequences (Fig 6A). One lineage (HTd) clustered with previously described elements from the ITmD37D family, also known as the *mat* family. The other two lineaged (HTb and HTf) are from the DD34D or *Mariner* family. Elements from the HTf network belongs to the *mauritiana* subfamily, and the HTb network elements, which are involved in most HTT events with species from outside of the Culicidae family, belong to the *irritans* subfamily. To obtain a more in-depth understanding of the HTb network, we performed a second BLAST search using as queries all sequences recovered from the search against non-mosquito genomes. We found HTb related elements to be present in 42 species of divergent invertebrates, including *W. bancrofti* (Fig 6B). Intragenomic dating of the horizontally transferred TEs provides more specific data and enables a more accurate interpretation of the HT events, including the source and sink species. The relative intragenomic age of TEs showed that the most ancient elements belong to the Lepidoptera species *Adoxophyes honmai*. Most of the flies’ elements are ancient and have similar ages. The elements of Asian mosquitoes (*An. maculatus, An. epiroticus, An. dirus, An. sinensis*) are more ancient than those of African ones (*An. gambiae, An. coluzzii, An. arabiensis, An. funestus*). Interestingly, the age of the *W. bancrofti* element is very similar to that of those of the Asian mosquitoes, suggesting that *W. bancrofti* acquired the element from the Asian *Anopheles* species and donated it to African species of *Anopheles* more recently (Fig 6C). Further evidence that this element is prone to transpose horizontally is provided by the multiple pairwise HTT significant signal among mosquitoes (Fig 6D).

**Fig 6:**
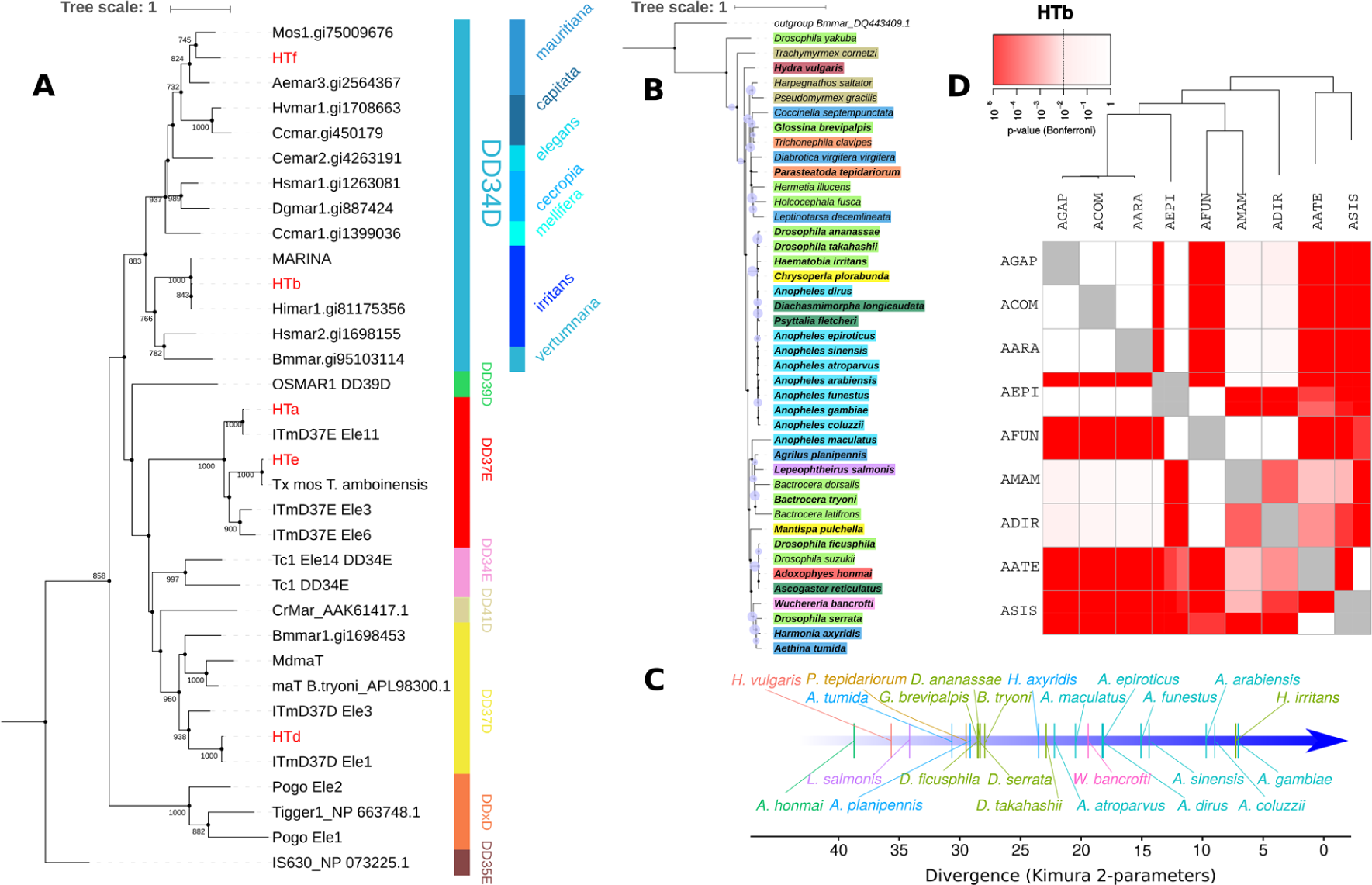
Transposons horizontally transmitted to other species. (A) A phylogenetic tree based on the transposase sequence of multiple ITm superfamily elements, including five Anophelinae TEs (labeled in red) involved in HTT events with other metazoan species and many elements that have already been described. The tree topology indicates that these five TEs belong to different families. HTb represents an element that is widespread in many metazoan genomes, as shown in part B. A species name in bold represents elements that share more than 80% of similarity to *Anopheles* ones. Brachycera is highlighted in light green, cnidarians in dark brown, ants in light brown, spiders in orange, beetle in blue, elements from the *Anopheles* genus in light blue, parasitoid wasps in dark green, Neuroptera in yellow, Lepidoptera in red, crustaceans in purple, nematodes in light purple. (C) A timeline built up using K2P distances showing the average relative age of elements in the genome of each species. (D) VHICA graphical representation of the HTb element in mosquitoes. The red-colored boxes represent statistically significant HTT signals.

## 3. Discussion

Most evolutionary biologists are still puzzled by the abundance and variability of the mobilome, even in closely related species. These rather simple sections of DNA are able to override the rules of Mendelian inheritance that govern the rest of the genome by two main ways: hijacking the host molecular machinery to generate more copies of themselves increasing in copy number while the majority of host genes remains stable and through horizontal transfer, a phenomenon that allows TEs to invade the genomes of other species despite being inherited vertically to all original host descendants [3,34]. More than hijacking the host molecular machinery, TEs also can be detrimental to their host due to transposition and recombination [4,35]. Nonetheless, host genomes possess a full arsenal of molecular mechanisms that can silence these elements [36–38]. Therefore, the abundance and diversity of the mobilome in the genome of each species is thus the result of an endless arms race between TEs and host [6,39,40]. Although much knowledge has been discovered about the TE “life cycle” since the genomic revolution there are still several open questions about a specific role of horizontal transfer in reshuffling TEs between different eukaryotic species and their impact on receptor species. In this study, we performed an in-depth mobilome characterization in all available mosquito genomes and detected a very large and variable TE content that is constantly exchanged by HTs between mosquitoes and distantly related species. Moreover, we found strong evidence of an intermediate worm species that mediated HT between mosquitoes and showed that horizontally transferred TEs significantly impacted the mosquito genome size.

The 24 mosquito genomes available have been scarcely studied from the mobilome perspective, except for the two model species *An. gambiae* and *Ae. aegypti* [22,41]. However, even though some TE sequences can be found in databases there is no standard annotation and the use of different methodologies for TE detection hinders a broad-scale comparative genomic analysis of the mosquito mobilome. Recently, 16 new *Anopheles* genomes have been published [25], but very little is known about the mobilome of these mosquitoes and annotated TEs have not been made available to the scientific community. To improve detection and standardized classification of TEs, we used a range of software that perform de novo TE detection and homology-based search followed by three different classification programs, PASTEC, TEsorter and RepeatClassifier. In line with other studies, the different methodological approaches generate different and yet complementary results [42,43]. On the one hand, *de novo* approaches result in more extensive TE content characterization in the case of most mosquito genomes. On the other hand, this method does not recover the whole mobilome. Some low-copy and more divergent TE families were recovered only by homology search. The most striking impact of using complementary methodologies in terms of overall TE content occurred in the case of the *Cu. quinquefasciatus* genome. Arensburger et al. (2010) showed that 29% of the *Cu. quinquefasciatus* genome was composed of TEs. A further study using a *de novo* method based on a structural search found other TEs not described at the time in this genome [44]. Here, we describe entire TE superfamilies that had not yet been described in some mosquito species and further expand the TE annotation for *Cu. quinquefasciatus* reaching a TE content of 43.55%. We also report for the first time the occurrence of Penelope elements in the genome of *Cu. Quinquefasciatus* and Crypton-like elements in the *Ae. albopictus* genome. Thus, a consistent characterization of the TE content of a given species requires that both *de novo* and homology-based approaches (with different reference TE database) be used for TE identification and classification.

Our in-depth mobilome characterization allowed us to investigate the influence of TEs on the size of the mosquito genomes. We observed that the mosquito genome fraction covered by TE correlates significantly with genome size although it is much more pronounced in species from the *Aedes* genus. Our results corroborate findings in arthropods [45] and vertebrates [46], However, it is interesting that the *An*. gambiae species complex showed a higher genome size to TEs ratio than the other *Anopheles* species. This suggests that other sequences are also responsible for genomic expansion in these species such as microsatellites and segmental duplications as shown for other species [47,48].

Given the high variability and uneven distribution of various TE superfamilies/families found in the genomes of mosquitoes, we next investigated the role of HTT in the evolutionary history of the mosquito mobilome. More than five thousand HTTs have been reported among eukaryotes so far, but only 4 HTT events involving mosquitoes have been well-documented, while 257 cases have been reported among drosophilids—a sister taxon [31]. Here, we report a fifty-fold increase (212) in the number of HTT events described among mosquitoes and investigate several hypotheses regarding the influence of host and TEs biological features on HTTs. One of the basic requirements for a given TE transfer to a new host species is the spatiotemporal overlap of the hosts [16]. Recent findings on Insects supported a higher HTT rate between taxa that shared the same realm [15]. However, in our fine-grained analysis of mosquito genomes, we found no association of ancestral species habitat range and HTT rate (S7 Fig) showing that different patterns may emerge driving HTT rate at different host taxonomic levels. Association between TE types or classes and the horizontal transfer rate have been linked showing that LTR and DNA transposons transfer much more frequently than non-LTR retrotransposons [34,49]. In mosquitoes, we found that half of the superfamilies that have undergone HTT belong to the LTR order of TEs, with an almost equal number of families of TIR and non-LTR elements in the remaining fraction. This result differs somewhat from other studies of insects investigating a broader set of taxa and a narrower time frame (last 10 MYA), which report a majority DNA transposons TEs involved in HTT events followed by LRT and non-LTR retrotransposons [15].

Another host intrinsic feature that may impact HTT rate is the phylogenetic relatedness of the host species. A number of studies have reported that horizontal transfer occurs more frequently in more closely related species [15,50]. However, long-range HTTs, between distantly related taxa, have also been observed [51,52]. These occur frequently with specific TE groups, such as Class II elements from the Tc1-Mariner [53] and hAT superfamilies [54]. Our data corroborate these findings, showing that most HTTs of Retrotransposons occurred between closely related species of the same genus, and long-range transfers among species of *Aedes* and *Anopheles* genus occurred mostly with DNA transposons TEs. Five TEs that underwent transfer to even more distant metazoan species were also Class II TEs from the Tc1-mariner superfamily. A similar pattern was observed in a study that investigated HTT in 195 insect species [15]. In general, HTTs occur more frequently between closely related species, but Class II TEs are more prone to long-range transfers probably due to host factor transposition independency and promoters that make them able to transpose in a wide set of divergent host species [55].

An interesting aspect of the large number of HTT events found among mosquitoes is that it is heterogeneous distributed on the mosquito evolutionary tree, showing a much higher prevalence of HT between species from the *Aedes* genus (Figure 2). Although there is a clear bias of more genomes available from *Anopheles* genus, we still found much more HTTs between *Aedes* species, the largest mosquito genomes sequenced so far and not a single HT involving *Culex quinquefasciatus*, a species with intermediate genome size (600Mb) but with a large TE content (around 40%). More unbiased studies including a diverse set of mosquito species should be performed to test these patterns, but our findings highlight that some taxa are more prone to exchange TEs horizontally than others, which is corroborated by other large-scale studies [56].

One of the most debated topics concerning HTT is how exactly do TEs move from one species to another and which vector species bridge the gap between species and facilitate HTT. Several long-range HTT events across phyla involve a Tc1-Mariner element of the *irritans* subfamily (Fig 6B). Interestingly, we found a number of copies of these elements in four different genome assemblies of *W. bancrofti—*a parasitic nematode transmitted by mosquitoes that causes lymphatic filariasis in humans [57]. At least one of these assemblies was reconstructed using only DNA sequences from adult worms isolated from the blood of an infected human patient (GCA_001555675.1), excluding any possibility of mosquito DNA contamination. The detection of copies of the element in independently sequenced *W. bancrofti* strains confirms that these elements are true components of the genome of this species. The *W. bancrofti* element has a similar age to that of those found in *Anopheles* species from Europe and Asia, although this element is younger in African *Anopheles* species. This suggests that *W. bancrofti* acquired this element from European and Asian *Anopheles* species and transferred them, more recently, to African *Anopheles*. Considering that *Anopheles* species do not migrate long distances, it is possible that these intercontinental transfers of TEs occurred by way of worms hosted by migrating animals, such as birds [58]. In addition, there is evidence showing that *W. bancrofti* is transmitted by different species of the *Aedes, Culex, Mansonia*, and *Anopheles* genera [59], suggesting that *W. bancrofti* likely mediated the transfer of HTb elements among at least eight species of *Anopheles* genus. The discovery of this parasitic nematode strengthens the evidence of parasitic species as an important agent of TE transfer among multicellular eukaryotic species.

Measuring the direct influence of horizontally transferred TEs is a difficult task since precise gene annotation of all genomes is required to associate TE insertions with the impact on surrounding genes. However, it is possible to estimate the proportion of the host genome that is derived from horizontally transferred TEs. This can be used as a proxy for estimating the likely impact/burden on the host genome. We found that horizontally transferred TEs contribute to genome size in varying degrees, depending on the mosquito genome. For both species of the *Aedes* genus, HT derived TEs represent a substantial portion of the genome (5.77-6.23%), that is, around 110Mb in the *Ae. albopictus* genome. This is similar to the *Drosophila melanogaster* genome size (143.7Mb) [60] and larger than that of the two-spotted mite *Tetranychus urticae* (91Mb) [61]. These results show that horizontally transferred TEs reach a high copy number since they replicate unchecked by the host genome after their invasion and can contribute substantially to the expansion and structure of the host genome. In line with that, genome size is also correlated with the proportion of the ge nome covered by horizontally transferred TEs, confirming that HTT contributed substantially to the mosquito genome size (Figure 2C).

Our study shows that complementary methodologies should be used for the precise characterization of the mobilome of a host species. Besides, our main findings are: mosquito mobilome varies in several orders of magnitude and is highly diverse; more than 212 TE families underwent horizontal transfer but we found no association between mosquito species spatiotemporal overlap and HTT rate which differ from other large-scale studies on Insects; there is a significant influence of the mobilome and horizontally transferred TEs on the size of mosquito genomes; several new long-range HTs involving mosquitoes and distantly related metazoans were characterized and among those we found a clear example of an intermediate species transmitted by mosquitoes enabling HTTs between various *Anopheles* species. Therefore, mosquito TEs are being constantly reshuffled among mosquito species by way of multiple horizontal transfer events facilitated by mosquito parasites such as the *W. bancrofti* roundworm.

## 4. Materials and Methods

### 4.1 Genomes Analyzed

All the mosquito genomes used in this study were obtained from VectorBase [62] and NCBI Assembly [63]. The GenBank accession codes are listed in S1 Table. The Mitochondrial DNA (mtDNA) was assembled using the raw reads for some species on Mitobin version 1.9.1 [64]. These sequences, along with mtDNA genomes already available in the NCBI, were used to build up a phylogenetic tree of studied species. The tree was constructed by way of maximum likelihood using PhyML [65] with 1000 bootstrap replicates, according to the GTR +G+I model based on the Akaike Information Criterion on SMS software [66].

### 4.2 Transposable element identification

Two main types of approaches are used to identify TEs in genomic assembled sequences: homology-based and *de novo* approaches. The most commonly used method involves the detection of homology between previously characterized TEs, normally obtained from databases such as Repbase, and the newly assembled sequences [30]. RepeatMasker is the software of the first choice for providing an overview of the TEs in any given genome [67]. Alternatively, *de novo* approaches identify TEs by their structural features or by repetitive/multiple-copy characteristics in the host genome [68–70]. There are various programs developed over the years, implementing these kinds of approaches, such as RECON [71], RepeatScout [72], PILER [73], LTR Finder [70], RepearExplorer [74] and so forth. As a way of avoiding underestimation, REPET pipeline [75] comes close to proposing a full pipeline for TE identification and annotation. Although each approach has its strengths, the use of either one of these methodologies in isolation almost always leads to an underestimation of the true TE content and adoption of a complementary strategy is therefore advised [42].

### 4.3 Identification of transposable elements using a *de novo* approach

A *de novo* approach was employed to identify mosquito TEs, using the TEdenovo pipeline from the REPET package [75]. See Supplementary Methods for details of pipeline execution. To complement our initial identification, we also ran RepeatScout, another software that uses a *de novo* approach [72], on the genome of each species using the default parameters. The repeated elements identified were then passed to the TEdenovo pipeline.

Given the virtual absence of elements characterized by TEdenovo or RepeatScout in some genomes, we examined the raw reads of these species further using Tedna [68], to ascertain whether TEs could have been removed before or during the genome assembly step.

### 4.4 Identification of transposable elements using a homology-based approach

At this stage, the TEs were identified based on homology with previously described TEs in the RepBase and TEfam databases. Initially, we executed a blastn and a tblastn search of the two databases, individually, against the genome of each species. Only HSPs (high-scoring segment pairs) with a bit-score above 200 were retained, to exclude random hits. Thereafter, blastn and tblastn-derived hits were merged into a single file to reconstruct the copies derived from HSPs that matched elements from the same family spaced no more than 1000 nucleotides apart.

All TE copies in a given genome were clustered by the CD-HIT-Est [76] algorithm with 80% identity and coverage using a global alignment strategy. Copies of each cluster were then extracted and aligned using the MAP algorithm, which reconstructed the representative consensus of each structural variant. Finally, a file containing all the TE consensuses for each species was generated for each of the databases used.

### 4.5 Consensus mapping and calculation of TE family divergence

The consensus sequences derived either from TEdenovo and homology-based search were classified using three different programs: PASTEC [77], a component of the REPET package; TEsorter, a recently described program that classifies TEs according to their conserved protein domains [78] and RepeatClassifier, a component of the RepeatModeler[79]. The consensus was then processed using RepeatMasker, which obtained the number of copies and base pairs covered by each TE of each genome. In the final TE dataset of each mosquito, we kept only TE consensus that masks regions of mosquito genome (Available in Figshare repository - https://figshare.com/s/1ea991f1a2004c3f8fa0). This is needed to remove false-positive TE consensus. We also extracted the Kimura 2-parameter distance (K2P) [80] from each TE family using RepeatMasker’s auxiliary scripts. This distance can be used to estimate the intragenomic age of these elements.

We performed an analysis of the correlation between the fraction of transposons and the genome size of each species studied. This was achieved using the cor() function (method = “spearman”) in the R software. The correlation coefficient was used to test the strength of the correlation. Resulting graphics were created using the ggpubr package [81].

### 4.6 Horizontal Transfer Analysis

To determine the inheritance mode of the TEs characterized, either vertical or horizontal, we used the R software’s VHICA package [82]. This package compares the relation between dS ratio and codon usage bias (CUB) linear regression of vertically-inherited single-copy orthologous host genes with transposable elements. A vertical transfer is the most likely scenario if the dS-CUB of a TE is not significantly different from that of the host genes. By contrast, a significant deviation in the host genes’ dS-CUB values indicates horizontal transfer.

Although the species that make up the *An. gambiae* complex are morphologically indistinguishable, we decided to investigate each one separately, in view of the many natural pre-mating barriers that restrict species hybridization. Furthermore, when mating does occur, there is evidence that male progeny are non-viable or sterile [83,84].

Fifty randomly selected single-copy orthologous genes from mosquito genomes were obtained from OrthoDB [85]. The ID of each of these genes was used to retrieve the nucleotide coding sequence for each gene from VectorBase (S4 File). The sequences for each ortholog gene set were then codon-aligned using the MACSE software [86].

The TE sequence consensuses of all species recovered using TEdenovo, excluding chimeric elements, were submitted to clustering by the CD-HIT-est algorithm, with 80% identity and 80% coverage using the refinement parameter (-g 1) and global alignment. As a validation method, we extracted the copy that was most similar to each consensus (based on the best bit-score hit) by conducting a BLAST search for each consensus against the genome of its respective species. This created a second set of clusters with the same structure as the consensus clusters, using the TE copies instead of the TE consensus. Clusters of representative TE copies remaining in the analysis were as follows: i) those having sequences in at least two species; ii) those in which all sequences had at least 600 nucleotides; iii) those having at least one sequence with ORF codifying a polypeptide greater than 200 amino acids in size. We used MACSE software to perform codon alignment for each cluster/family (as defined by Wicker et al.), taking the nucleotide sequence of the largest ORF found among the clustered sequences as a reference (-seq parameter) and the remaining sequences as a FASTA file in the -seq_lr parameter. The flanking regions of sequences, based on the beginning and end of the reference sequence after alignment, were trimmed, as only the coding region was of interest for the dS-CUB analysis.

The alignment of the orthologous genes and TEs were then passed as input to the VHICA package to ascertain whether vertical/horizontal transfers occurred. Those TE clusters whose p-value was less than 0.01 in a one-tailed statistical test were considered horizontal transfer events. Additionally, the percentage of the genome involved in horizontal transfer events, as well as the K2P parameter for each TE family in each species, was calculated using the RepeatMasker software and its auxiliary scripts.

To investigate whether any of the TE families that were involved in horizontal transfer events among the 24 studied mosquito species could also be involved in horizontal transfers to other species, we performed a blastn (dc-megablast) of the sequences of these TEs against: the NCBI nt database; all genomes of protostomes, plants, fungi, protists, flatworms, viruses, Echinodermata, Hemichordata, and Chordata organisms present in NCBI as of January 2019. Those matches that had a high degree of identity at the nucleotide level, with more than 80% of the mosquito’s TE sequence coverage, and those that had a copy number greater than five were considered to be probable HTTs. When a species had more than one assembly from different samples, we dispensed with the need for a copy number if the matches were present in the majority of the different genome assemblies of this species.

### 4.7 Dating Horizontal Transfers

The dating of HTT events was performed by applying the formula T = k/2r [87], where **T** is time, **k** is the synonym substitution rate (dS) between TE copies from two species, and **r** is the evolutionary rate of the species groups. We obtained dS estimates per mosquito taxon from the 50 single-copy ortholog genes as follows: 17.567 × 10^−3^ mutations per million years for transfers among mosquitoes from the *Anophelinae* subfamily, 9.205 × 10^−3^ for transfers within Culicinae subfamily of mosquitoes, and 10.006 × 10^−3^ for transfers between the Anophelinae and Culicinae subfamilies (S1 File). It should be noted that we found very few orthologous TE copies and these were restricted to species of the *An. gambiae* complex. As the speciation time is not well defined among these species, we performed HTT dating using host gene estimates.

### 4.8 Analysis of the geographical distribution of vector mosquitoes

The distribution region of each mosquito species of the genus *Anopheles* was taken from the distributions presented by Sinka et al. [88–91] and the distribution predicted using the *Malaria Atlas Project* [92]. These regions were considered the ancestral habitats of these mosquitoes since most *Anopheles* mosquitoes are not invasive species and thus disperse very little. We consider sub-Saharan Africa to be the ancestral habitat of *Ae. aegypti* [93] and the east and southeast Asian region extending to India to be the native habitat of *Ae. albopictus* [94]. To evaluate whether the overlap and/or proximity of the mosquitoes species distribution has any impact on the likelihood of the horizontal transfer, a point-biserial correlation analysis was performed using the cor.test () function of the R software, considering the number of transfers that occurred between two species of mosquitoes and their geographical overlap or non-overlapping distribution.

## Acknowledgments

We are thankful to the Bioinformatics Center of the Aggeu Magalhães Institute.

## Supporting Information Captions

**S1 File. Supplementary methods**.

**S2 File. VHICA output images**. Figures of VHICA graphs for each case of horizontal transposon transfer detected.

**S3 File. Similarity matrix of Tc1-mariner**. Pairwise similarity matrix of five TE families that are horizontally transferred among several of 24 studied species to different species.

**S4 File. Coding region of orthologous genes**. Nucleotide sequence of coding regions used for VHICA analyses.

**S5 File. Accession numbers and name of species that are searched for HHT by blast**. The genome IDs, taxa, and species/virus names of more than 32,000 genomes that were searched for horizontally transferred mosquito TEs.

**S1 Table. List of mosquito genome assemblies used in this study**.

**S1 Fig – Fraction of genome occupied by TEs characterized by both methods, *de novo* and homology-based**.

**S2 Fig – Mobilome fraction of each Wicher’s system Orders**.

**S3 Fig – Intragenomic dating of the four most abundant TE Orders of mosquito genomes**.

**S4 Fig. P-values distribution of all positive pairwise comparisons**. The majority of detected HTT signals are very significant.

**S5 Fig. Landscape of TE families that undergone horizontal transfers**. Each dot represents the relative age of each family in a mosquito genome.

**S6 Fig. Horizontal transfer of a TE of R4 clade of R2 superfamily**. Each red square represents a significant HTT pairwise comparison.

**S7 Fig. Lack of correlation between geographical overlap and the number of horizontal transfer cases in mosquitoes**. Each dot represents the number of horizontal transfers between to species.

